# Layer-specific dynamics of local field potentials in monkey V1 during electrical stimulation

**DOI:** 10.1101/2024.10.19.619242

**Authors:** Sangjun Lee, Zhihe Zhao, Ivan Alekseichuk, Sina Shirinpour, Gary Linn, Charles E. Schroeder, Arnaud Y. Falchier, Alexander Opitz

## Abstract

The mammalian neocortex, organized into six cellular layers or laminae, forms a cortical network within layers. Layer specific computations are crucial for sensory processing of visual stimuli within primary visual cortex. Laminar recordings of local field potentials (LFPs) are a powerful tool to study neural activity within cortical layers. Electric brain stimulation is widely used in basic neuroscience and in a large range of clinical applications. However, the layer-specific effects of electric stimulation on LFPs remain unclear. To address this gap, we conducted laminar LFP recordings of the primary visual cortex in monkeys while presenting a flash visual stimulus. Simultaneously, we applied a low frequency sinusoidal current to the occipital lobe with offset frequency to the flash stimulus repetition rate. We analyzed the modulation of visual-evoked potentials with respect to the applied phase of the electric stimulation. Our results reveal that only the deeper layers, but not the superficial layers, show phase-dependent changes in LFP components with respect to the applied current. Employing a cortical column model, we demonstrate that these in vivo observations can be explained by phase-dependent changes in the driving force within neurons of deeper layers. Our findings offer crucial insight into the selective modulation of cortical layers through electric stimulation, thus advancing approaches for more targeted neuromodulation.

## 1. Introduction

The mammalian neocortex, cytologically characterized by its six distinct layers, forms an interconnected network crucial for processing cortical information ^1, 2^. In the sensory cortex, neurons in layer 4 mostly receive feedforward inputs from the thalamus, which are then strongly projected to layers 2/3 for further processing ^3^. Subsequently, these inputs are forwarded to layers 5/6, where recurrent inputs to layer 2/3 originate ^4^. Local field potentials (LFPs) have been recorded from those layers using multisite laminar probes to understand the microcircuit of cortical columns in sensory cortex ^5, 6, 7^. Especially, the primary visual cortex (V1) has been widely explored due to its well-characterized connectivity and distinct cell types in each layer ^8, 9, 10^. Moreover, layer-specific LFPs can be effectively investigated by applying a visual stimulus, which serves as a sensory evocation to generate feedforward activation of the canonical cortical circuit ^11, 12, 13^. LFP activity reveals that an early current source occurs in the input layer (layer 4), followed by current sink in the superficial layers (layers 1-3) and deep layers (layers 5/6).

Sensory-evoked LFPs show distinct spatiotemporal patterns across cortical layers. One notable observation is that evoked LFPs in deeper layers are stronger than those in superficial layers ^6^. A previous study demonstrated that flash-evoked LFPs increase with more depth in the rat brain in response to visual stimuli ^14^. This pattern emerges because sensory input is primarily injected into layer 4, as well as due to cytological structure differences among the layers. For instance, large pyramidal cells in layer 5 generate strong dipoles along the dendrites, contributing to larger LFPs ^14^. In addition, the variation in firing rates across layers affects LFP amplitude, as LFPs reflect the summation of synaptic activity from neural spikes in neuron populations ^14^. Cortical column modeling of mouse V1 has shown that neurons in layers 2/3 have the lowest firing rates, whilst those in layers 5/6 have the highest firing rates ^15^. As a result, when thalamic input generated by sensory stimuli is injected into the model, layers 5/6 exhibit larger LFPs ^16^.

Electrical stimulation has emerged as a promising therapeutic application for treating various neurological and psychiatric symptoms such as depression, epilepsy, schizophrenia, and Parkinson’s disease ^17, 18, 19, 20^. However, it is still unknown how electrical stimulation modulates evoked LFPs across cortical layers. While there is a comprehensive understanding of how the phase and amplitude of electrical stimulation affects neural dynamics at the level of single neurons ^21, 22, 23^, the extension of this knowledge to cortical layers requires careful investigation. How electrical stimulation affects layer-specific LFPs is crucial for bridging the gap between individual neural activity and the synchronized response of neural populations to external currents across layers. This understanding is also important for translating insights from layer-specific LFPs to human EEG, which represents the cumulative activity of LFPs ^14^. To achieve this, we recorded LFPs while applying electrical stimulation in nonhuman primates (NHPs). NHPs have emerged as significant models for studying the biophysical and physiological effects of electric stimulation ^21, 24, 25, 26, 27, 28, 29^, at various spatial scales, due to the similarity of their cortical layers to those of humans ^30^.

Here, we record LFPs using laminar probes across all layers of V1 in two lightly anesthetized nonhuman primates (NHPs), employing a flash visual stimulus to evoke sensory responses while simultaneously applying 1.5 Hz alternating current (AC) transcranially to the occipital lobe. We first suppress stimulation artifacts while preserving LFPs using an independent component analysis (ICA) algorithm. We demonstrate that electrical stimulation can selectively modulate LFP activity in deeper cortical layers (layers 4–6), by changing the amplitude of LFPs in a phase-dependent manner. We observe that the LFP components, the first positive peak (P1) and negative peak (N1), show preferential increases depending on the AC phase, highlighting the phase-dependent nature of LFP modulation. Furthermore, we use a cortical column model of V1 to investigate the layer-specific effects observed in vivo experiments. Our findings suggest that distinct neural activity of pyramidal neurons in deeper layers contributes to an enhanced response to external currents. Our results provide valuable insights into the casual relationship between neural activity within cortical layers and the LFPs in response to electrical stimulation.

## Results

Monkeys (*n* = 2) were surgically implanted with a multisite probe with 23 contacts in the V1 while they sat on a primate chair, receiving flash visual stimuli generated by a light stimulator (Figures 1A and 1C). The probe was inserted perpendicular to the cortical surface, considering the alignment of the pyramidal neurons. Layers were identified by analyzing the distribution of current source density and electric field (Figure 1B; see Method). Two sets of LFPs were recorded: one without electrical stimulation (Flash condition) and another with electrical stimulation (Flash + AC condition). The aim was to examine how electrical stimulation affects the modulation of LFPs in a layer-specific manner. We removed the AC artifacts in laminar recordings using the ICA algorithm^31^ while preserving the visual evoked LFPs (Figure 1D). In the Flash + AC condition, the power spectrum density showed a complete rejection of the AC frequency (1.5 Hz) compared to the power of electrical stimulation (Figure 1F).

**Figure 1.**
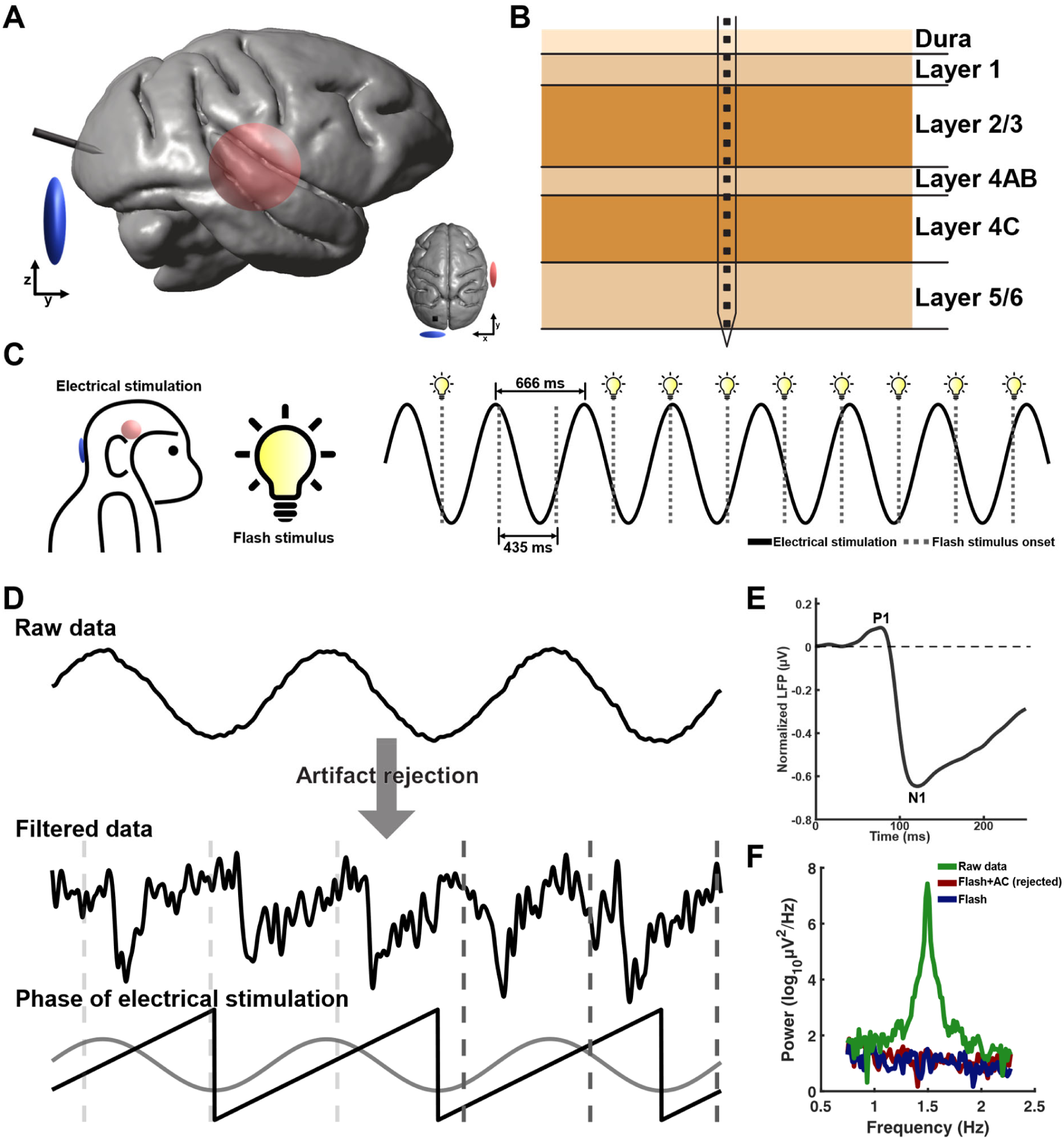
Experimental design for obtaining visual-evoked LFPs in V1 layers during 1.5 Hz alternating current. A) A multisite laminar probe was inserted into the monkey’s V1. One electrical stimulation electrode (blue) was positioned on the scalp near the probe, while the other one (red) was placed over the right temporal area. B) Schematic illustration showing the placement of the contacts across cortical layers in V1. C) Visual stimuli were delivered to the monkeys at a frequency of 2.3 Hz using a high-intensity light stimulator. Concurrently, electric current oscillating at 1.5 Hz was injected via the electrodes. D) Raw LFPs were processed to remove AC artifacts while preserving visual-evoked LFPs. 1.5 Hz AC signal was extracted by applying a bandpass filter between 0.5 Hz and 2 Hz. The phase of AC was calculated using the Hilbert transformation for further phase dependency analysis. E) Illustration of visual-evoked LFP, with the first positive peak labeled as P1 (approximately 75 ms from the visual onset) and the first negative peak labeled as N1 (approximately 120 ms). Power spectrum density demonstrates that the algorithm for the artifact removal effectively eliminates AC artifacts in the Flash + AC condition.

### Effects of electrical stimulation on LFP modulation in layer-specific manner

Visual-evoked LFPs were recorded across contacts and normalized relative to the largest amplitude of LFPs in layers 5/6. LFPs exhibit higher amplitude in the deeper layers (layers 4–6) compared to the superficial layers (layers 1–3) in both monkeys (Figure 2A and Figure S1B). The increased strength of LFP amplitudes at deeper depths may be attributed to the higher synaptic and neural activity occurring in deeper layers. These findings are in line with previous studies that have recorded LFPs in the sensory cortex ^32, 33^. Notably, the amplitude of LFPs after 100 ms from the onset of visual stimulus was higher under the Flash + AC condition compared to the Flash condition, especially in the deeper layers. This difference is evident in the changes in LFP components, P1 and N1, across layers (Figure S1C). For monkey 1, the mean N1 amplitudes in layers 5/6 were 0.59 ± 0.1 (Flash + ES condition) and 0.47 ± 0.12 (Flash condition), while the mean P1 amplitudes were 0.13 ± 0.08 and 0.12 ± 0.09, respectively. A similar trend was observed in monkey 2, with the mean N1 amplitudes of 0.38 ± 0.12 with AC and 0.28 ± 0.06 without AC, and the P1 amplitudes of 0.10 ± 0.09 and 0.06 ± 0.06, respectively.

**Figure 2.**
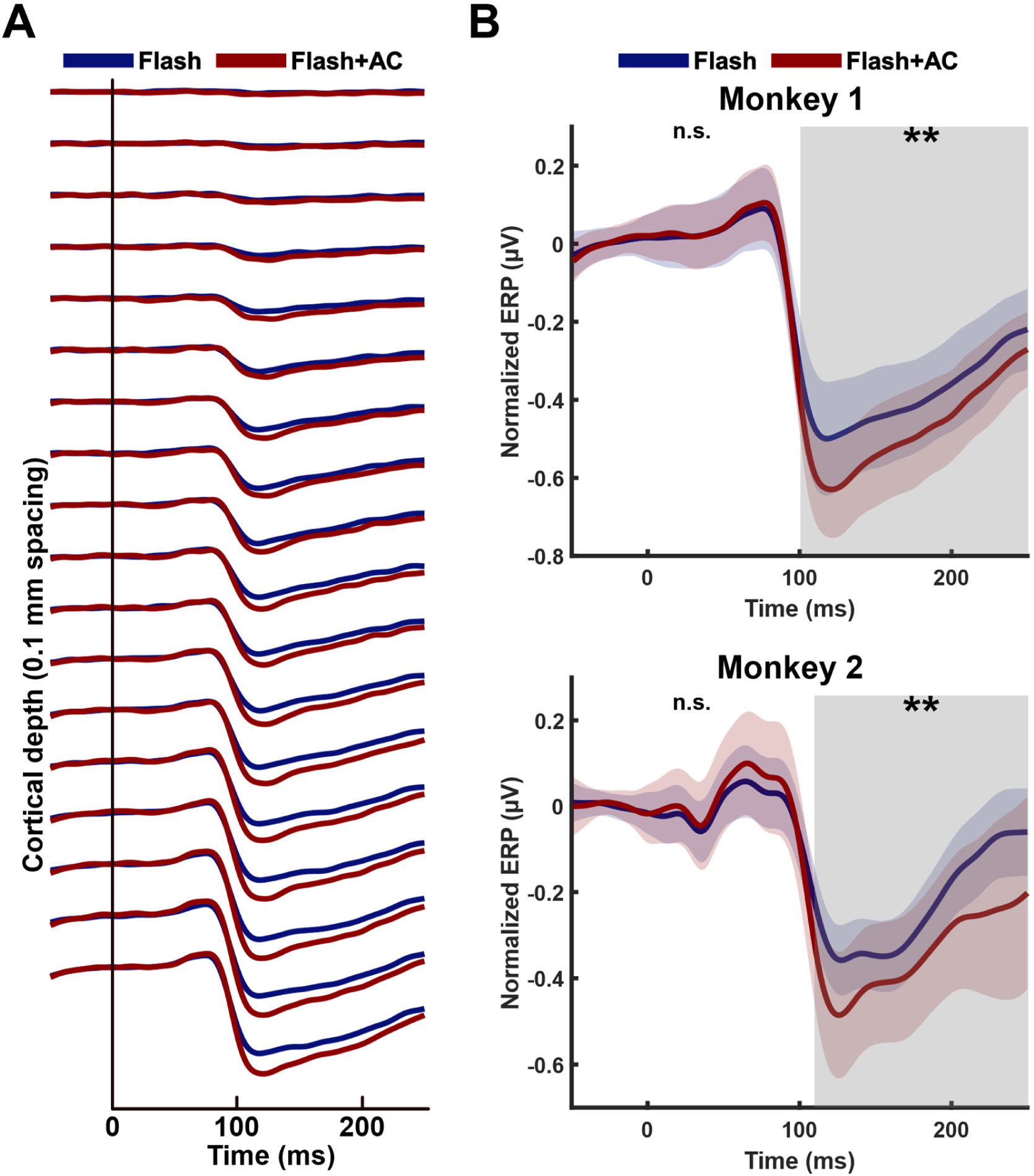
Effects of electrical stimulation on layer-specific visual-evoked LFPs in V1. A) Local field potentials (LFPs) recorded using multisite probe for monkey 1. LFPs along contacts (0.1 mm spacing) during the Flash condition (blue line) and the Flash + AC condition (red line). LFPs were normalized relative to the last contact, which has the largest LFP, followed by averaging them across trials at each contact. Time indicates the duration from the flash visual stimulus onset. B) LFPs in layers 5/6 for both monkeys 1 and 2. Normalized LFPs were averaged across trials and contacts within layers 5/6. Thick lines and shades represent the averaged LFP and standard deviation, respectively. The cluster-based permutation test determined significant differences between the Flash and Flash + AC conditions across contacts in the time range from 0 ms to 250 ms. Significant differences occurred within the contacts in layers 4–6 after 100 ms from the onset of the flash visual stimulus. The gray shade represents time windows where significant differences were observed (\*\**p* < 0.01; n.s., not significant).

We performed a cluster-based permutation test to test for the statistical significance between Flash and Flash + AC conditions. We found a significant difference between the two conditions across contacts ranging from layer 4AB to layers 5/6 in the time range between 100 ms and 250 ms for monkey 1 (*p* = 4.99 x 10^-4^) and monkey 2 (*p* = 9.99 x 10^-4^). No significant clusters were observed during the time range of 0 ms to 100 ms (Figure 2B and Figure S2). Unlike for monkey 1, in monkey 2 we observed a significant difference between the two conditions across contacts in layers 2/3 between 100 ms and 140 ms. This may be due to the lower volume conduction effect of LFPs from the input layer to layers 2/3 in monkey 2 during Flash condition ^34^. Our findings indicate that the amplitude of LFPs evoked by sensory stimuli increased after 100 ms, especially in the deeper layers (layers 4–6). This suggests that electrical stimulation enhances the amplitude of the N1 component.

### Phase dependency of layer-specific LFPs during electrical stimulation

Next, we performed a phase dependency analysis to investigate whether the amplitude of the LFP components, P1 and N1, changes depending on the phase of AC. The trial-based LFPs were sorted into 20 phase bins based on the phase of AC, which was determined by the phase at the onset of the visual stimulus. Then, we determined the P1 and N1 components from LFPs, followed by averaging them across trials for each layer for each phase bin. Our findings show that the P1 component was increased during specific phases of AC for deeper layers in monkey 1. The phase dependence of the P1 amplitude was strongly distributed in a bimodal circular pattern in layers 4–6 (Figure 3B). The circular distribution of the P1 amplitude showed bimodal mean directions across different layers: −145° for layer 1, −135° for layers 2/3, −114° for layer 4AB, −118° for layer 4C, and −113° for layers 5/6. The bimodality in the N1 amplitude was less pronounced as in P1. The mean directions of the bimodal distribution of the N1 amplitude were as follows: −40° for layer 1, −61° for layers 2/3, −41° for layer 4AB, −37° for layer 4C, and −40° for layers 5/6. Monkey 2 exhibited a unimodal circular distribution of the LFP components based on AC phases, with a similar trend as in monkey 1, showing a strong directionality for the P1 amplitude (Figure 3C). The unimodal mean direction of the P1 and N1 amplitude was −159° and 100° for layer 1, −86° and 30° for layers 2/3, −94° and 74° for layer 4AB, −59° and 107° for layer 4C, and −110° and 53° for layers 5/6.

**Figure 3.**
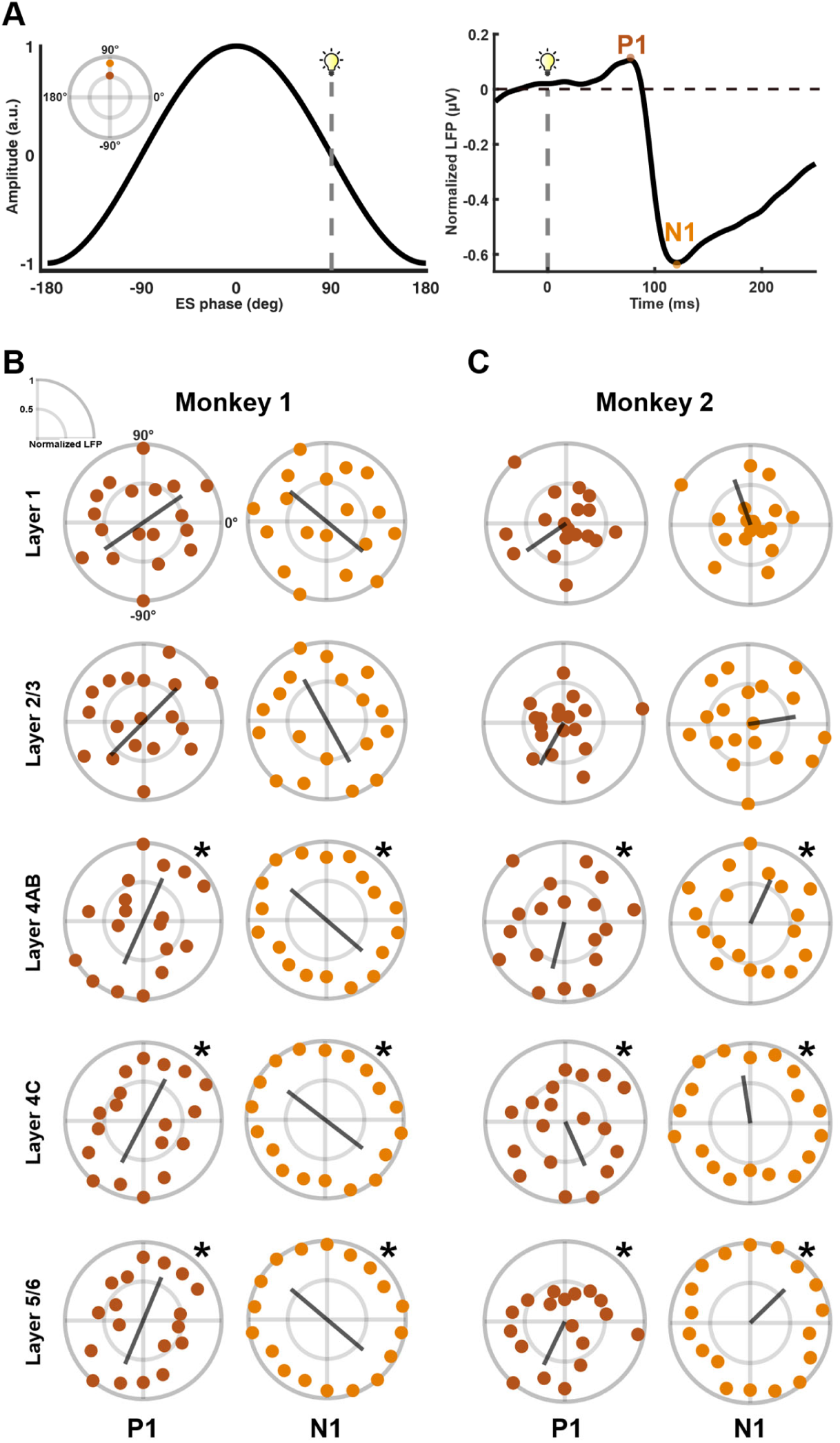
Layer-specific phase dependency of LFP components during electrical stimulation. Phase dependency of the amplitude of LFP components, P1 and N1 to external stimulation. The P1 and N1 components were sorted into 20 phase bins, followed by taking trial- and phase bin-averages for each layer. The gray lines represent the mean direction of the phase preference of LFP amplitudes. A) Illustration of the sorting of LFP components based on the AC phase. For instance, in a single trial, the visual stimulus was presented at a time corresponding to an AC phase of 90° (left). For each trial, the amplitudes of the P1 and N1 components were calculated and sorted into the phase bin that includes 90° (right). B) Circular distributions of P1 and N1 amplitudes according to the AC phase show a bimodal characteristic for monkey 1. C) Circular distributions show a unimodal characteristic for monkey 2. A permutation test revealed significant directional preferences in P1 and N1 amplitudes with respect to the phase of AC only within deeper layers (layers 4–6) (\**p* < 0.05) for both monkeys.

We performed a permutation test for each layer to investigate the significance of the amplitude changes in LFP components relative to the phase of AC. This was done by comparing the vector length in the mean direction of the original data to the vector lengths obtained from the permutations. The vector length quantifies the strength of phase-locking across AC phases ^35^. Our findings demonstrate that P1 and N1 amplitudes show a preferred directionality with respect to the phase of AC only in deeper layers (layers 4–6). No significant effects were observed in superficial layers (layers 1–3) for both monkeys (Figure S3). The amplitude of P1 in deeper layers was larger during the phase in which the amplitude of N1 was smaller for both monkeys. This raises the question of whether the observed phase dependency is caused by electrical stimulation or an inherent physiological phenomenon. For instance, it is possible that the onset of the visual stimulus happens to coincide with a specific bio-signal oscillating at 1.5 Hz. As a control analysis, we performed the same phase dependency calculation for the data from the Flash condition for a virtual AC frequency of 1.5 Hz. No significant phase preference of the amplitude of LFP in both P1 and N1 components were found across all layers (Figure S4). Thus, our results suggest that the amplitude of the LFP component is selectively modulated depending on the phase of an external current in a layer-specific manner.

### No LFP modulation in white matter

Additionally, we recorded LFPs in white matter in monkey 1 as a control region to investigate whether LFP modulation is restricted to deep cortical layers. To do this, the laminar probe was lowered so that several contacts of the probe were located within white matter. Then, the same analysis procedures used for the cortical layers were employed to determine P1 and N1 components, followed by sorting them into 20 phase bins. Our results show that electrical stimulation has a minimal effect on the modulation of LFPs in white matter, in contrast to its impact on cortical layers (Figure 4A). A cluster-based permutation test revealed no significant modulation of LFPs between the Flash and Flash + AC conditions. In the phase dependency analysis, the amplitudes of P1 and N1 showed no preferred directionality with regard to the phase of AC (Figure 4B). These findings suggest that electrical stimulation modulates LFP activity selectively within deep layers in the gray matter.

**Figure 4.**
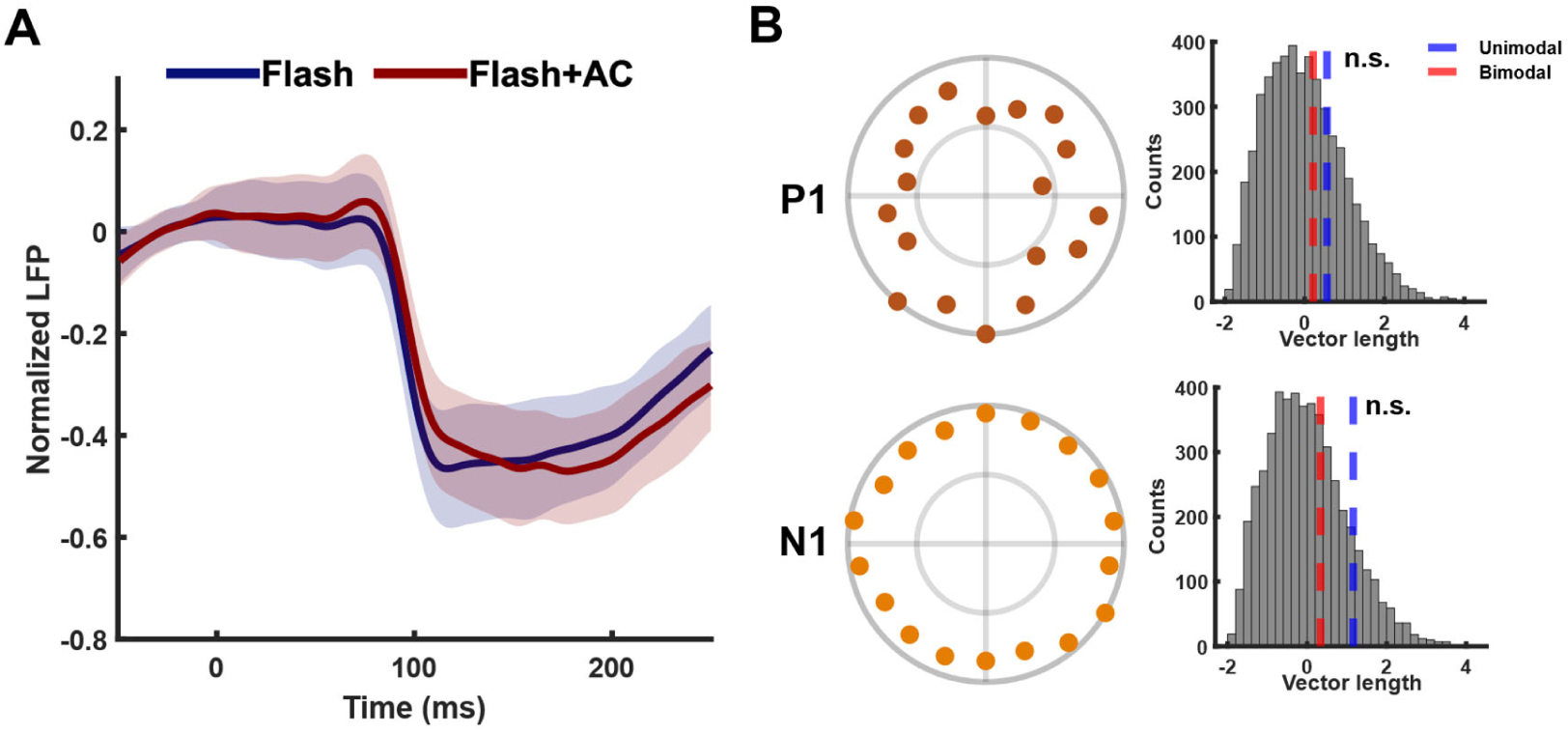
Effect of electrical stimulation on LFP activity in white matter. A) Local field potentials (LFPs) in white matter in monkey 1. Normalized LFPs were averaged across trials and contacts within white matter. Thick lines and shades represent the averaged LFP and standard deviation, respectively. Time indicates the duration from the flash visual stimulus onset. B) Circular distributions of the amplitude of LFP components, P1 and N1, depending on the phase of AC in white matter in monkey 1 (left column). The results from the permutation test are depicted for P1 and N1 components (right column). The permutation test shows that there is no significant directional preference in P1 and N1 amplitudes with respect to the phase of AC, regardless of whether the circular distribution was unimodal or bimodal (n.s., not significant). The gray histogram represents 5000 permuted vector lengths. The blue lines and red lines indicate the unimodal vector length calculated from the original data and the bimodal vector length, respectively.

### Biophysics of electrical stimulation across cortical layers

To investigate the relationship between LFP changes and the biophysics of electrical stimulation, we measured the electrical voltage and electric field across cortical layers (see Table. S1 in the Supplementary Material). In monkey 1, the electric field exhibited an initial increase from layer 1, reaching a considerable peak in layers 2/3, and thereafter decreased in deeper layers (Figure 5A). The average electric fields were as follows: 0.62 V/m in layer 1, 2.53 V/m in layers 2/3, 1.18 V/m in layer 4AB, 0.62 V/m in layer 4C, and 0.37 V/m in layers 5/6. Monkey 2 showed a similar electric field distribution across cortical layers. However, in contrast to monkey 1, higher electric fields were delivered to deeper layers (Figure 5B). The average electric field was to be 0.97 V/m in layer 1, 2.76 V/m in layers 2/3, 1.6 V/m in layer 4AB, 1.59 V/m in layer 4C, and 1.42 V/m in layers 5/6. Interestingly, despite the relatively high electric field in layers 2/3, the effect of electrical stimulation on the LFP modulation was not observed in these layers. This suggests that neurophysiological properties of cortical layers have a stronger effect on LFP modulation than the physics of electric stimulation.

**Figure 5.**
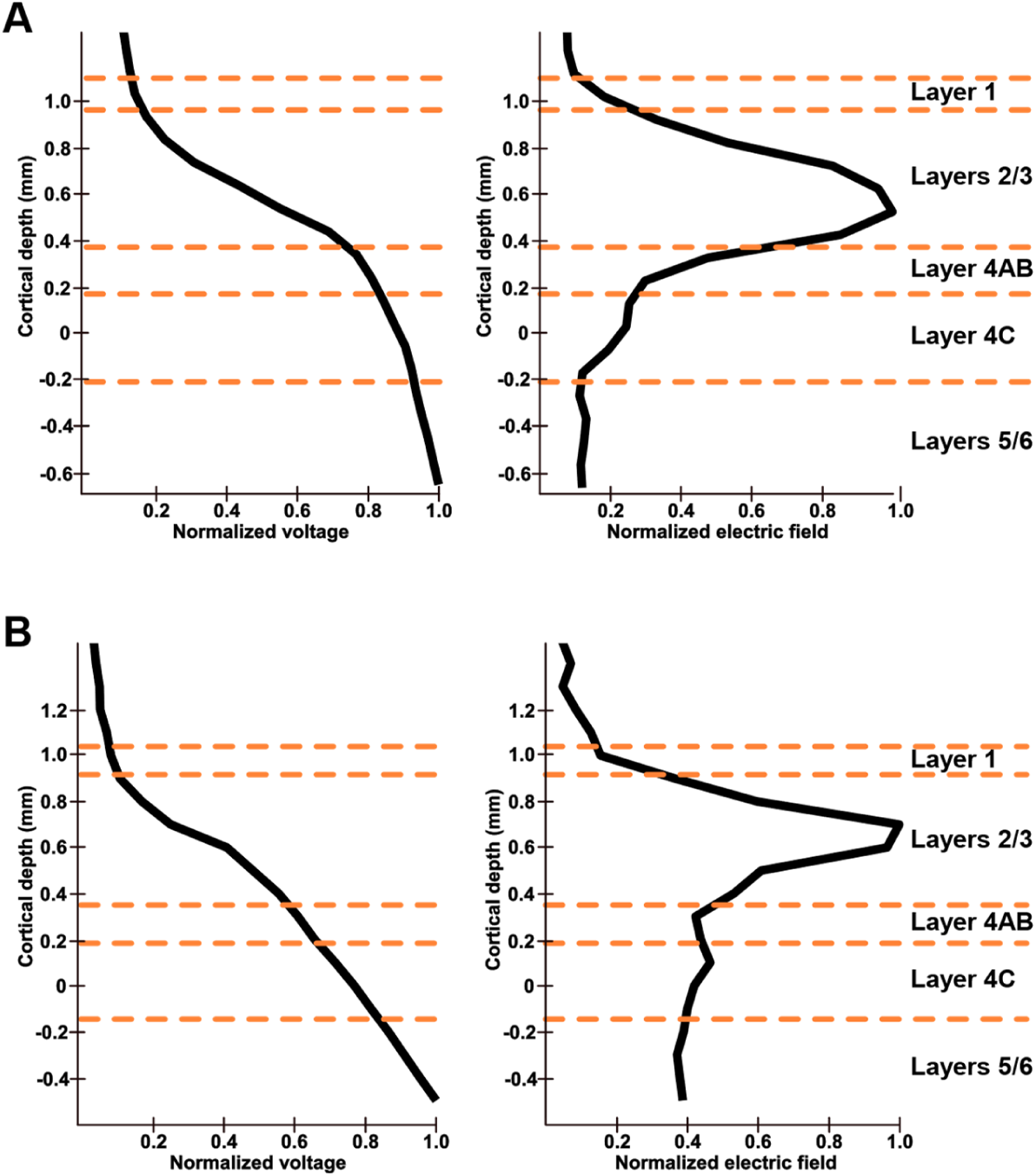
Biophysics of electrical stimulation in V1 layers. Distributions of electrical voltage and electric field across cortical layers in A) monkey 1 and B) monkey 2. The electrical voltage and electric field values were normalized to their maximum values. The electric field was calculated by taking the gradient of the voltage along the direction of the laminar probe. For both monkeys, the electric field began to increase from layer 1, reaching the maximum in layers 2/3, and then decreased in deeper layers.

### Neural activity in cortical column model

In order to further elucidate how observed layer specific changes in LFPs can be explained, we extended a cortical column model of V1^15^ to integrate AC stimulation. Before applying AC, we investigated how neural activity arising from the flash stimulus varies across layers. Our model results show that excitatory neurons in the deeper layers have a distinct firing response around 60 ms after the visual stimulus onset (Figure 6A). Enhanced firing rates are present in layers 5/6, followed by layer 4 and layers 2/3 (Figure 6B), which aligns with stronger LFPs in layers 5/6 (Figure S5A). Firing rates in layers 5/6 were determined by averaging the rates from both layers to align with in vivo findings, whereas layer 1 was excluded due to the absence of excitatory neurons.

**Figure 6.**
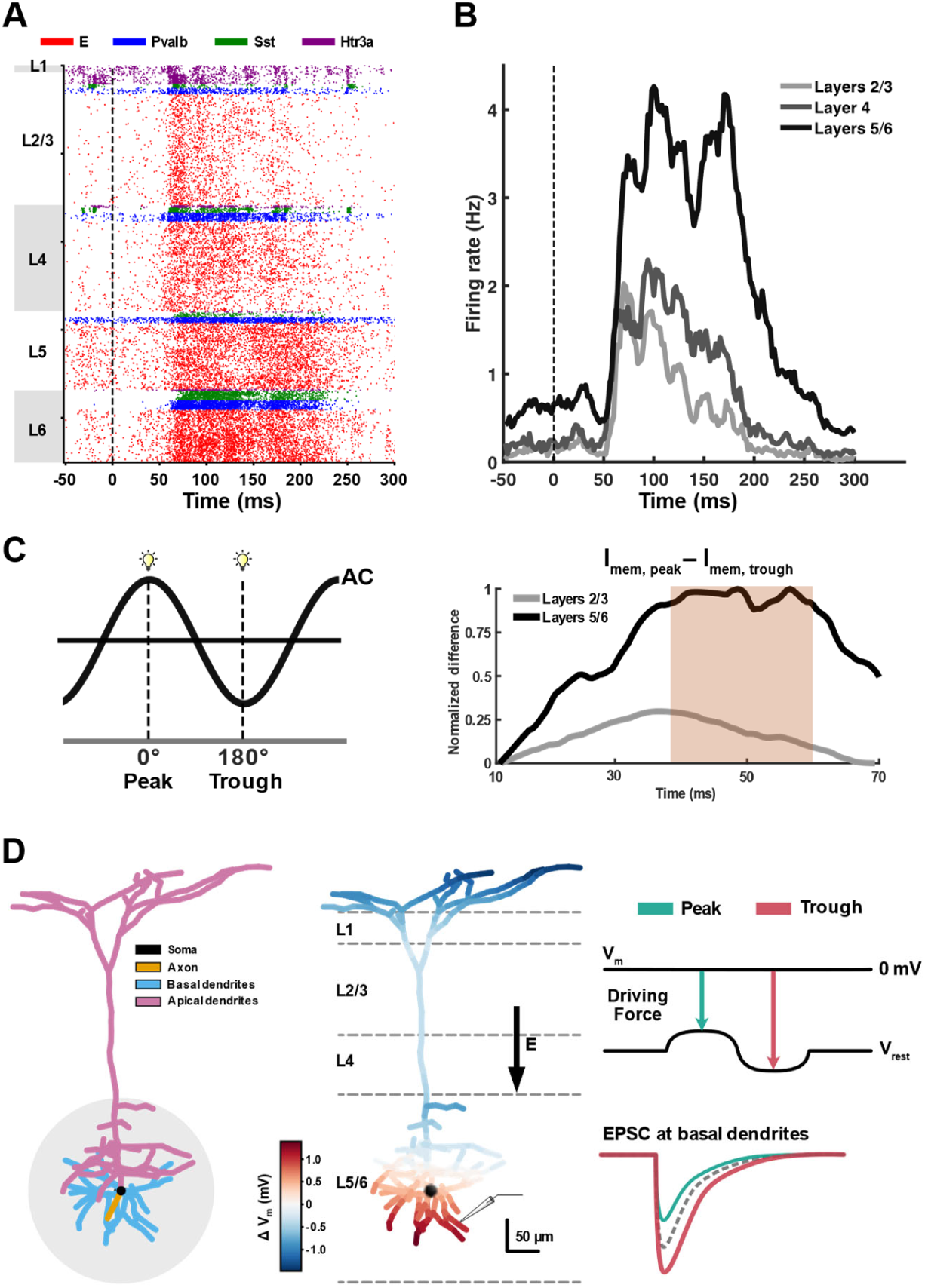
Neural activity in the cortical column model of V1 during flash stimuli. A) Raster plot showing the neural spiking of different types of neurons across layers. The red dots represent excitatory neurons, while the others represent parvalbumin-positive interneurons (blue), somatostatin-positive interneurons (green), and 5-HT3a receptor-positive interneurons (purple), respectively. B) Averaged firing rate of excitatory neurons across layers over time, calculated using a 2 ms time bin. Dash lines indicate the onset of flash stimuli. C) Membrane current from the basal dendrites of 500 neurons in both layers 2/3 and layers 5/6 was calculated under two conditions: when a flash stimulus was applied at the peak phase and the trough phase of AC (left). The difference between the membrane currents in the peak condition (I_mem, peak_) and trough condition (I_mem, trough_) is the highest during the period of P1 in LFP around 50 ms. The red shade represents the period of P1. D) Schematic illustration explaining the phase dependency. The synaptic input from lateral geniculate nucleus (LGN) enters the region adjacent to basal dendrites in deeper layers, which are highly responsive to this input (left). When the peak phase of AC flows downward direction (middle), the membrane potential in basal dendrites is depolarized, leading to a weaker driving force. It results in weaker (less negative) excitatory postsynaptic current (EPSC) affecting change in LFPs and vice versa during the trough phase.

Next, we applied AC in the model to investigate the layer-specific phase dependency observed in in vivo LFPs. Interestingly, our model showed that the P1 (about 50 ms) occurred before neural firing (Figures 6A and S5A), suggesting that factors other than neural firing may contribute to the strong phase preference of P1 in our observations. We hypothesized that the membrane current (I_mem_) induced by thalamic input around 50 ms (Figure S5C) in the basal dendrites, mostly located in layers 5/6, is modulated in a phase-dependent manner. Given that I_mem_ is influenced by the membrane potential, which is responsive to the phase of the AC, or the polarity of the electric field, these phase-dependent changes in I_mem_ likely contribute to the phase dependency of LFP. To test this, we calculated I_mem_, an indirect feature determining the LFP deflection, when a flash stimulus was applied at either the peak or trough phase of AC. Our results show that the difference in I_mem_ between peak and trough conditions was positive during the P1 period, with a stronger difference in the deeper layers (Figure 6C). This suggests that the peak phase of AC produces a more positive (or less negative) I_mem_, leading to a more positive LFP relative to the trough condition. This finding elucidates the distinct phase preference of P1 specifically in the deeper layers rather than the superficial layers, as shown in Figures 3B and 3C. On the other hand, the trough phase of AC induces a more negative I_mem_, resulting in a larger negative LFP compared to the peak condition. The membrane current during the peak phase consistently exceeds that in the trough phase. This results in opposite phase preferences for the positive and negative LFP deflections, as seen in our in vivo observations.

## Discussion

Electric current stimulation can alter the membrane potential of neurons, causing either depolarization or hyperpolarization ^23, 36, 37^. LFPs represent extracellular measurements of transmembrane currents arising from postsynaptic potentials from synchronized neuronal populations ^14, 38^. Thus, a change of membrane polarization from external electrical stimulation will affect LFPs. Our results demonstrate that visual-evoked LFPs during AC are enhanced compared to no stimulation, but only in deeper layers (layers 4–6). This indicates that neurons in deeper layers show a greater response to AC compared to those in superficial layers. This can be explained by the unique anatomical and physiological features of neurons in different layers. Layer 4 is characterized by its high neuron density ^16, 39^, and layer 5 includes larger pyramidal cells compared to layers 2/3 ^39, 40^. This can result in a stronger response to AC due to the increased number of neurons producing synchronized activity. In fact, deeper layers exhibit higher firing rates in response to visual stimuli compared to layers 2/3 ^41^, and layers 2/3 show reduced synchronization of neural populations compared to deeper layers ^42, 43^. This effect is also demonstrated in our cortical column model, showing that the neural firing rate in deeper layers is higher than in layers 2/3 (Figure 6B). In white matter, LFPs did not show a modulation of evoked potentials by electrical stimulation (Figure 4). The dominant components of the WM are the complex fiber tracts, resulting in insufficient neural activity for the generation of response to electrical stimulation ^44, 45^. This highlights the anatomical and functional specificity of different cortical layers in their responsiveness to electrical stimulation.

Interestingly, the physiological effects observed in our study cannot simply be explained by the biophysics of electrical stimulation. We found that the electric field strength in layers 2/3 was at least twice as strong as in layers 5/6 for both monkeys (Figure. 5). This suggests that layers 2/3 include certain anatomical features that reduce electrical conductivity, leading to higher electric fields. Lower conductivity increases the tissue’s resistance to current flow, causing a more concentrated and increased electric field. This reduced conductivity may be due to the high density of blood vessels and astrocytes ^46^, whose endothelial cells form gap junctions that raise electrical resistance ^47^. Indeed, computational simulations demonstrated that electric fields are markedly higher when the blood vessels are included in simulations during electrical stimulation ^48^. Despite the stronger fields, layers 2/3 did not show modulation or phase dependency in LFPs, unlike deeper layers. Therefore, we conclude that LFP modulation is more sensitive to layer-specific neural features than to the amplitude of the electric field.

A phase preference for the first LFP component P1 only emerges in deeper layers, suggesting that neurons in layers 5/6 have a stronger neural response to specific AC phases. One possible explanation for this observation lies in the modulation of the electrochemical driving force by AC phase-induced changes in membrane potential. These alternations in the driving force affect the postsynaptic currents ^49^. As the excitatory postsynaptic current (EPSC) changes, the resulting fluctuations in membrane currents lead to changes in the extracellular potential, as reflected in LFPs (Figure. 6D). We focused on basal dendrites because they are the primary sites for synaptic inputs and integration of neural activity ^50^. During the peak phase of AC, basal dendrites are depolarized, leading to a weaker driving force. Conversely, during the trough condition, they become hyperpolarized, resulting in a stronger driving force. This difference is evident in the membrane current, which is higher under the peak condition (Figure 6C). Thus, a weaker driving force produces a less negative ESPC, resulting in a more positive (less negative) LFP deflection. In our in vivo experiments, the P1 component had a preferred phase around −110° for both monkeys, corresponding to the rising phase of AC. During this phase, the driving force consistently increased throughout the period of the evoked LFP following the visual flash. This effect is layer-specific, as the basal dendrites of large pyramidal neurons in the deeper layer show a high response to AC, which can spread to adjacent layer 4 due to volume conduction ^51^.

Our explanation can elucidate why the P1 and N1 components have distinct preferred phases. Unlike the P1, the N1 peak appears to be more associated with neuronal firing, as it occurs after the spiking of pyramidal neurons in V1. Indeed, it shows a strong correlation with the firing rate as depth increases (Figure S5B). Neurons in V1 become less excitable during the preferred phase when the P1 peak is stronger, likely due to a weak driving force. This results in a lower N1 amplitude since neurons are less likely to fire during this phase. On the contrary, when the P1 peak is lower, neurons are more excitable, which leads to an increased N1 amplitude.

Our results come with some limitations. In our experiments the monkeys were lightly anesthetized to minimize interference from other neural activities. Stimulation under awake conditions might differ, however sensory-evoked LFPs during anesthesia are well-preserved compared to the awake state ^52^. To improve separability of AC stimulation from ongoing LFPs, we used low-frequency stimulation to minimize the potential influence of LFP modulation caused by the entrainment of intrinsic oscillations, such as alpha, beta, and gamma rhythms ^53, 54^. Our results depend on the successful rejection of tACS artifacts in the LFPs. Although residual artifacts could have an influence, we consider it unlikely due to the consistent outcomes across cortical layers and the absence of effects in white matter. Additional recording modalities such as single-unit activity could further strengthen our results. We observed some differences in LFP modulation between the two monkeys. This might originate from differences in the electric field strength across the layers. In monkey 2, the electric field in deeper layers was somewhat higher than in monkey 1, resulting in a more pronounced increase in LFP and displaying a strong unidirectional phase preference in LFP modulation. Our group has previously demonstrated dose-dependent neural entrainment during tACS in resting-state NHPs ^22^. Thus, future work might profit from recording dose-responses relationships to investigate the relationship between current intensity and LFP modulation.

In conclusion, our findings demonstrate that sensory-evoked LFPs are modulated by electrical stimulation in a layer-specific and phase-dependent manner. This modulation was predominantly observed in deeper layers, where large pyramidal neurons are more responsive to external electric fields. Our cortical column model suggests that changes in the driving force in deeper layers can explain the observed phase dependency. This study advances our understanding of how electric fields interact with cortical microcircuits and highlights the influence of layer specific properties on LFP modulation. These insights can inform future strategies to optimize stimulation parameters, targeting specific layers to improve the therapeutic efficacy of neuromodulation.

## Methods

### Subjects

Two female capuchin monkeys (*Cebus apella*), each aged 15 years and weighing between 1.5 and 3 kg, were included in the current study. All procedures were approved by the Institutional Animal Care and Use Committee of the Nathan Kline Institute for Psychiatric Research. Surgical procedures corresponded to the methods that were previously established ^55^. Under general anesthesia, we conducted craniotomy. The monkeys were implanted with a headpost, and a Cilux recording chamber (Crist Instruments) positioned over the occipital cortex to aim electrode penetrations perpendicular to the primary visual cortex (Area V1). Both implants were fastened using MRI compatible ceramic screws (Thomas Recording, GmbH). For each recording session, we lowered a 23 channels linear array electrode (U-Probe, Plexon) which has been used in previous studies ^32, 56, 57^ through a grid to ensure a perpendicular insertion into area V1. The electrode intercontact spacing was 0.1 mm and the impedance range for the contacts was 300-750 kΩ. We lowered the probe until the contacts would bracket all cortical layers, as well as the area outside the cortex. We confirmed that the probe covered all cortical layers using current source density (CSD) analysis, as described in the later method part. The LFPs outside the cortex (i.e., dura mater) were necessary to establish the border of layer 1 during electrical stimulation. We verified that the electric field did not dramatically change within the area outside of the cortex, while it began to increase noticeably in layer 1.

### Visual stimuli

Flash visual stimuli were used to induce sensory-evoked LFPs in the cortical layers. The visual stimuli were presented using a Grass strobe light photo stimulator while the monkey was seated in a primate chair, with its head fixed, in an electrically shielded chamber under dim lighting conditions. The monkeys were lightly anesthetized with 1–1.5% isoflurane. The distance between the monitor and the monkey’s eyes was 86 cm. Visual stimuli (*n* = 200 to 400) were presented at a frequency of 2.3 Hz, meaning that a visual stimulus appeared every 435 ms. The monkeys underwent two experimental sessions: first, a session without electrical stimulation (Flash condition), and then a session with electrical stimulation (Flash + AC condition). To mitigate the visual fatigue, we ensured ample rest between the two sessions.

### Laminar recording

Two monkeys underwent laminar recordings, where a multisite probe was implanted into their V1 region, for both Flash and Flash + AC conditions. Electrical signals were recorded at 23 contacts within the probe, with an intercontact spacing of 0.1 mm. The impedance ranged from 0.3 to 0.5 megohms. The signals were then amplified by a factor of 10 and filtered (bandpass DC of 10 kHz) using the preamplifier from Plexon. To obtain LFP activities, the signals were analog filtered within the range of 0.1 Hz to 500 Hz, followed by downsampling to 2 kHz.

### Electrical stimulation

Electrical stimulation was transcranially employed using a Starstim system (Neuroelectrics, Barcelona, Spain) through Ag/AgCl electrodes attached to the scalp. One electrode was placed over the occipital lobe, right under the location where laminar probes are inserted. Another electrode was attached to the right temporal area (Figure 1A). The electrodes were round, with a radius of 10 mm. The electrode montage was chosen to ensure that the electric field is more likely to be directed parallel to the laminar probes. We preliminary calculated the maximum electric field during the experiment and found that the maximum electric field in monkey 2 was lower than that in monkey 2 when injecting the same current into both monkeys. Therefore, we decided to inject the intensity of the current (peak-to-zero) of 0.1 mA for monkey 1 and a higher current of 0.2 mA for monkey 2. The frequency of the current was chosen to be 1.5 Hz to allow for the generation of various AC phase values corresponding to the onset of the visual stimulus presented at a rate of 2.3 Hz (Figure 1C).

### Data preprocessing

The data were analyzed using Matlab 2022b (MathWorks). LFPs were obtained by applying a 4th-order Butterworth bandpass filter between 0.5 Hz and 100 Hz to the raw signals ^58^, followed by downsampling to 1 kHz using the Fieldtrip toolbox ^59^. We chose to apply a cut-off frequency of 100 Hz to remove possible contamination from multiunit spiking responses in the higher frequency range, up to 300 Hz ^60^. Additionally, the notch filter was employed to eliminate the powerline noise at 60 Hz. LFPs were segmented into 300 ms epochs ranging from −50 ms to 250 ms relative to the onset of the flash visual stimulation. Then, baseline correction was employed using the time window from −50 ms to 0 ms for every epoch. For the condition with electrical stimulation (Flash + AC condition), the AC artifact-filtered data, as described in the following section, was used to segment epochs. In each trial, the most dominant positive peak that appeared about 75 ms after the onset of the visual stimulus was identified as P1 within the time range of 55–90 ms, while the negative peak occurring around 120 ms, within the time window of 100–140 ms, was labeled as N1 for all laminar probes (Figure 1E). Trials that exceeded the mean ± 5 standard deviation (SD) were regarded as artifacts and thus excluded from further analysis ^55^. For monkey 1, there were a total of 320 trials in the Flash condition and 216 trials in the Flash + AC condition, whereas for monkey 2, there were 395 trials in the Flash condition and 216 trials in the Flash + AC condition.

### Artifact rejection of electrical stimulation

The AC artifact rejection was performed on the data after lowpass and notch filtering, before segmentation, using Matlab 2022b (MathWorks). Independent component analysis (ICA), a method to statistically decompose mixed signals into independent sources, has been widely employed to remove the artifact induced by AC ^61^. In the current study, we employed the second-order blind identification (SOBI) algorithm ^62^, a type of ICA, embedded in the Fieldtrip toolbox ^59^, to remove the AC components oscillating at a sinusoidal frequency of 1.5 Hz ^31^. After applying SOBI, sinusoidal components oscillating at 1.5 Hz were detected using the Fourier fast transformation (FFT). Although we attempted to detect harmonics, such as 3 Hz and 4.5 Hz, within the components, they were not found. This might be due to the direct measurement of signals in laminar recordings, which are minimally affected by volume conduction, unlike electroencephalogram (EEG) signals recorded on the scalp ^63^. Then, we employed a discrete Fourier transform (DFT) filter to subtract the component from the data ^64^. This approach has been widely used to reject the powerline noise oscillating in a sinusoidal waveform, allowing us to specifically reject the 1.5 Hz component in our study. Finally, the components are backprojected to the contact level, resulting in LFPs that were cleansed of AC artifacts. Power spectrum analysis was then conducted to evaluate the effectiveness of AC artifact rejection. As shown in Figure 1F, AC artifacts are successfully removed from the data in Flash + AC condition.

### Biophysics of electrical stimulation

The data were analyzed using Matlab 2022b (MathWorks) and the Fieldtrip toolbox. To obtain electric potential oscillating at 1.5 Hz, the raw data from laminar recordings was filtered using bandpass filtering from 0.5 Hz to 2 Hz with a fourth-order zero-phase, forward-reverse Butterworth filter, followed by downsampling to 1 kHz. Given that the signals were amplified by a factor of 10, we divided the electric potential by 10 to determine the actual amplitude of electrical stimulation. Then, the electric fields were calculated by applying the numerical gradient to the measured electric potential along contacts ^27^. As the laminar probes are aligned in the radial direction with the cortical surface, we were able to quantify the electric field in the radial direction at each contact. Using FFT, we extracted the phase and amplitude of electric fields at each contact for the maximum frequency ^26^, which is the frequency of AC oscillating at 1.5 Hz (Figure 1D). For each trial, the phase of AC at the onset of the visual stimulation was determined for each contact to identify the specific phase at which the visual-evoked LFP occurred. This enabled us to assign the phase value of AC for each trial.

### Layer-specific LFP analysis

Laminar contacts were assigned to layers based on three approaches: i) current source density (CSD), ii) electric field distribution, and iii) anatomy. The CSD was computed by taking the second derivative of visual-evoked LFPs (filtered and segmented) along the contacts, using the data from Flash condition ^65^. CSD analysis is capable of capturing the early feedforward input in layer 4C during visual stimulation ^32^. Based on the current sink/source ^12^ and anatomical references ^10, 30^, the contacts were assigned to a specific layer. However, pinpointing the precise location where layer 1 begins in the CSD distribution is challenging. Thus, we defined layer 1 as the region where the electric field begins to increase, marking the boundary between the cortex and the outside of the cortex (Figure 5). Subsequently, the contacts were categorized into five layers: four contacts were assigned to layer 4C and layer 5/6, two contacts to layer 4AB, five or six contacts to layer 2/3, and one or two contacts to layer 1 for both monkeys. Then, we obtained the layer-averaged visual-evoked LFPs for each trial for each layer. Furthermore, the laminar contacts were lowered to capture LFP activity in the white matter for the control condition. We lowered 1.3 mm from the base location to ensure that the five contacts from the tip of the probe were positioned within the white matter, rather than in the cortical layers. Then, the same procedure was repeated with the five contacts to obtain LFPs in the white matter.

### Phase dependency analysis

Phase dependency analysis was conducted using the data from the condition with electrical stimulation (Flash + AC condition) to investigate whether LFP activity varies according to the phase of AC across cortical layers. Data were analyzed using Matlab 2022b (MathWorks) and CircStat toolbox ^66^. In the previous step, the phase of AC was labeled for each trial. Hence, the layer-averaged trials were able to be sorted into 20 phase bins of 18° for each layer. The LFP components, P1 and N1, were then determined for each phase bin, followed by averaging the trials to obtain the mean value of the LFP components. Last, we computed the mean direction and vector length of the phase-binned P1 and N1 components for each layer. The mean direction represents the preferred AC phase for which the LFP component tends to be strong, and the vector length indicates how strongly LFP components preferentially align with the AC phase. Interestingly, we found a strong bimodal circular distribution of LFP components according to the AC phases in monkey 1, while monkey 2 showed a unimodal distribution. For instance, in monkey 1, the P1 component showed higher amplitudes at AC phases of 90° and −90° simultaneously, resulting in convergence in mean direction to 0°, and vector length to zero. To address this issue in monkey 1, we employed a different approach that can calculate the mean direction and vector length in the bimodal circular data by using the function *circ_axialmean* embedded in the CircStat toolbox ^67, 68^.

### V1 cortical circuit model

We utilized the biophysically detailed cortical column model of V1 established in the previous study ^15^. This model incorporates both multicompartment and leaky-integrate-and-fire (LIF) neurons, with a total of 230,924 neurons arranged in a cylindrical structure. It includes 17 neuronal classes distributed across cortical layers, with distinct excitatory and inhibitory populations. Thalamocortical input is provided by a lateral geniculate nucleus (LGN) module, while background activity is simulated using a Poisson process. This comprehensive model accurately represents the cortical circuitry and connectivity patterns of V1. The stimulus used in the simulation consisted of full-field flashes. Each trial comprised a 500 ms gray screen period, followed by a 50 ms white screen, and then a 350 ms gray screen. Ten trials were simulated.

The LFP evoked by the flash stimulus in simulations was derived from the extracellular potential using a fifth-order Butterworth low-pass filter with a cutoff frequency of 100 Hz, followed by a downsampling to 1 kHz, and baseline correction (−50 ms to 0 ms). LGN spike trains used as input to the V1 circuit were generated with the FilterNet module provided with the model, using 17,400 ‘LGN units’ ^15^. The original model’s synaptic arrangement was preserved: LGN-to-excitatory V1 neuron synapses were located on dendrites within 150 μm of the soma, while LGN-to-inhibitory V1 neuron synapses were placed on the soma and dendrites without distance constraints. The background input, firing at 1 kHz with a Poisson distribution, was not stimulus dependent. The recurrent connection probabilities and synaptic placements from the original model remained unchanged in this study. Then, firing rates were calculated for each layer using a 2 ms time bin.

To couple the electric fields to the V1 cortical circuit model, electric potentials were calculated for each neuron segment based on the external electric field along the vertical axis of the model and were set as the extracellular potentials in the ‘extracellular mechanism’ within the NEURON environment ^69^. Given that an electric field is considered quasistatic, it can be divided into spatial and temporal components ^70^. We employed a sinusoidal 1.5 Hz tACS waveform with simulated electric field strengths of 16 mV/mm, matching experimental values, based on the previous simulation results ^71^. The tACS was initiated at 250 ms and sustained for the remainder of the simulation. To investigate the phase dependency of the visual evoked response, the tACS waveform was shifted to align specific phases (peak or trough) with the flash onset at 500 ms. To assess the effects of LGN synaptic input on LFP, we recorded membrane currents in passive basal dendrites of 500 randomly selected excitatory neurons per layer for several key reasons. First, LGN synaptic inputs primarily target dendrites within 150 μm of the soma. Second, tACS-induced depolarization/hyperpolarization effects are substantially stronger in large, elongated excitatory neurons. Third, excitatory neurons constitute approximately 85% of the neuronal population in each layer, contributing more significantly to LFP. Measurements were taken for two phase conditions and a control condition (tACS with silent network). We summed currents across all segments and subtracted the control condition currents from the phase conditions to isolate LGN-induced effects from tACS-induced fluctuations.

### Statistical analysis

All statistical analyses were conducted using Matlab 2022b (MathWorks). For the comparison between Flash and Flash + AC conditions, we employed a nonparametric cluster-based permutation test to investigate whether electrical stimulation significantly improves visual evoked LFPs along the contacts ^59, 72, 73^. This approach is especially advantageous for comparing the data in time series, as it provides robust findings without excessively conservative thresholds, such as those in Bonferroni correction ^72^. Concisely, the trials from Flash and Flash + AC conditions were compared using a *t*-test for each contact and time point and combined as a cluster according to their temporal adjacency. Then, cluster-level statistics were calculated by taking the sum of the *t*-value within the cluster. We set the permutation iteration to 2000 times and a critical value of 0.01 to strongly prevent false positives. Given that the contacts were assigned to specific cortical layers, we can easily determine which layers exhibit a significant effect on LFP modulation by an external current.

The permutation test was also performed to evaluate the significance of the change in the amplitude of P1 and N1 depending on AC phases ^74^. For each component, the labeled phase was randomly shuffled to generate surrogate data, followed by sorting it into the phase bin and calculating the mean direction and vector length as in the previous step. Given that the phase-binned surrogate data did not have bimodality, we employed unimodal circular calculations in the permutation test. Shuffling was repeated 5000 times, resulting in 5000 values for the vector length. We standardized the vector length from the original data to a z-score using the mean and standard deviation from the surrogate distribution. Next, a one-tailed p-value was calculated to determine the proportion of surrogate data that was smaller than the original data. If the original data exceeded 95% of the surrogate values, it was considered statistically significant. The same process was employed for each layer.

## Acknowledgments

This research was supported in part by RF1MH124909, R01EB034143 and the University of Minnesota’s MnDRIVE Initiative.

## Data and code availability

The datasets generated and analyzed during the current study are available from the corresponding authors on reasonable requests.

## Author Contributions

SL, AYF, CS and AO conceived the study and designed the experiments. ZZ conducted neuron modeling. IA and SS advised on data analysis. GL, AO, and AYF acquired and preprocessed the data. AO supervised the study.

## Supplemental information

Document S1. Figures S1–S5 and Table S1.

