## Supplementary Materials for "Layer-specific dynamics of local field potentials in monkey V1 during electrical stimulation"

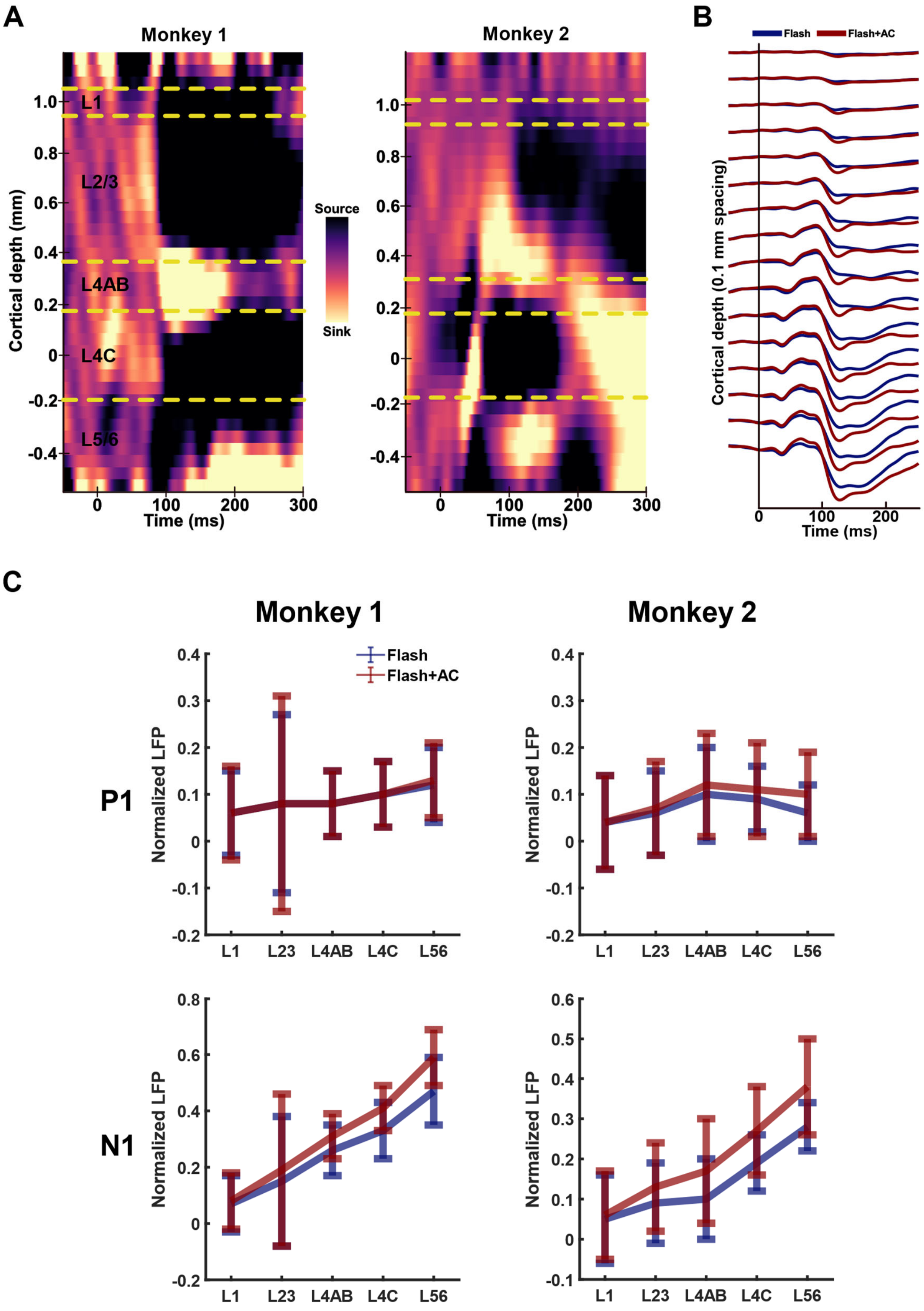


**Figure S1.** A) Depth profile of the current source density (CSD) signals averaged across trials. The color yellow represents the current sink, while the color black represents the current source. B) LFPs recorded using multisite probe in monkey 2. Only the contacts involved in cortical layers are illustrated. Red lines represent visual-evoked LFPs in the Flash + AC condition, and blue lines depict LFPs in the Flash condition. C) Comparison of layer-averaged amplitudes of LFP components, P1 and N1, between the Flash (blue) and Flash + AC (red) conditions for both monkeys.

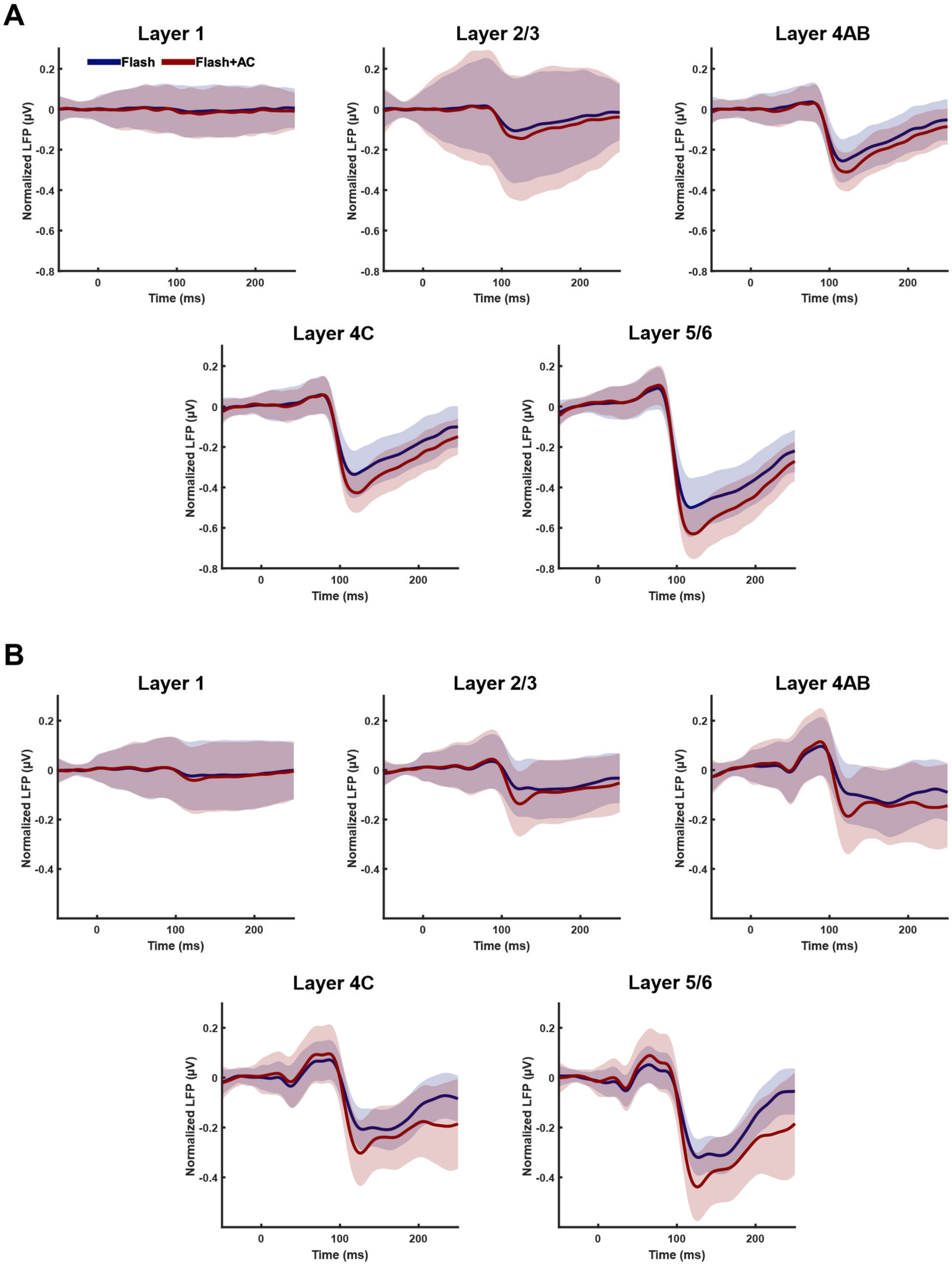


**Figure S2.** Effects of AC on layer-specific visual-evoked LFPs in V1 for A) monkey 1 and B) monkey 2. Normalized LFPs were averaged across the trials and contacts within each layer. Thick lines represent averaged LFP in the Flash condition (blue) and Flash + AC condition (red), with shades representing the standard deviation.

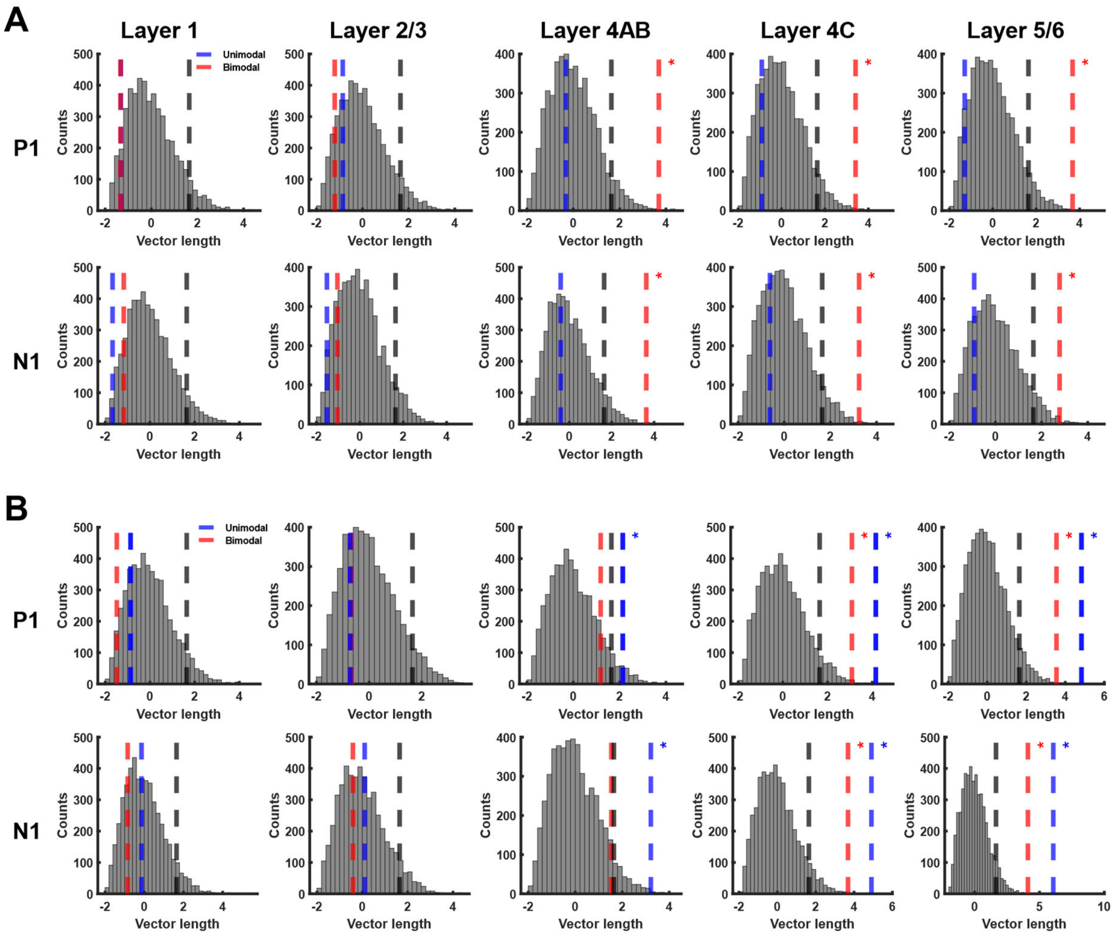


**Figure S3.** Permutations test in the phase-dependent analysis corresponding to Figure. 3 for A) monkey 1 and B) monkey 2. The gray histogram represents 5000 permuted vector lengths. Gray dotted lines indicate the significance level. The blue and red lines indicate the unimodal vector length and bimodal vector length obtained from the original data, respectively. For monkey 1, the permutation test reveals significant bimodal phase preference in P1 and N1 with respect to the phase of AC within deeper layers (layers 4–6). For monkey 2, there is a unimodal phase preference in the amplitude of LFP components depending on the phase of AC in the deeper layers.

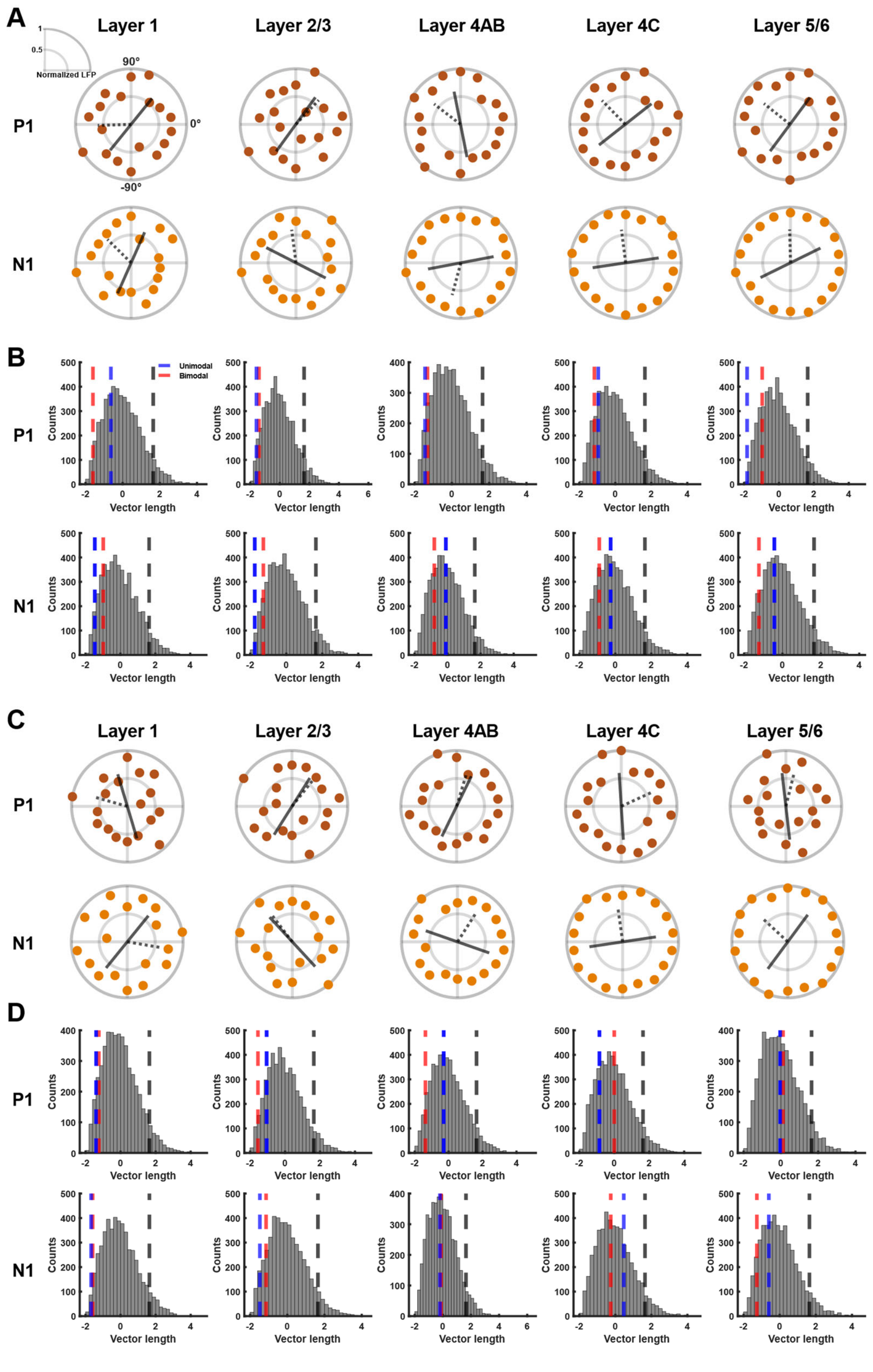


**Figure S4.** Amplitude of P1 and N1 components according to the phase of virtual AC in the Flash condition for both monkey 1 (A and B) and monkey 2 (C and D). A, C) The P1 and N1 components were sorted into 20 phase bins, followed by taking trial- and phase-averages for each layer. Gray thick and dotted lines represent the bimodal mean direction and unimodal mean direction of the amplitude of LFP components, respectively, based on the phase of virtual ES. B, D) Permutation test shows the absence of significant directional preferences in LFP component with respect to the phase of AC across all cortical layers when virtual AC is assumed to be applied in the Flash condition.

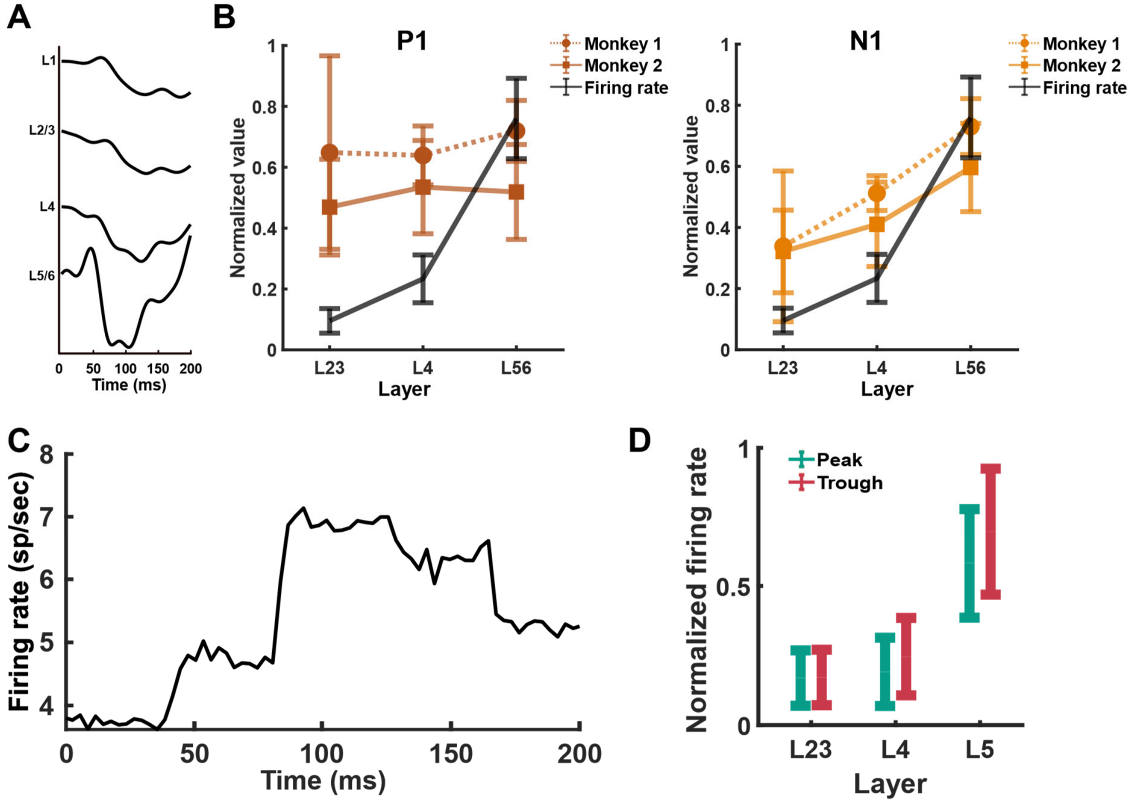


**Figure S5.** A) LFPs evoked by the flash stimulus in the cortical column model across layers. B) Comparison of LFP components (P1 and N1) obtained in in vivo experiments and the firing rate between 50 ms and 150 ms. The P1 component does not correlate with an increase in the firing rate (left), while the N1 component shows a pattern that corresponds more strongly with the increasing firing rate as depth increases in both monkeys (right). C) Firing rate of LGN neurons, showing an initial spike around 50 ms (corresponding to the P1 period in simulations), followed by a second firing arising around the period of neural firing of V1 neurons. D) Normalized firing rate of V1 neurons when the visual stimulus was applied at the either peak or trough phase of AC. The firing rate is higher during the trough phase of AC than the peak phase in the deeper layers, while there is no comparable difference in firing rate between two phase conditions in the superficial layer.

### Supplementary Table

**Table S1.** Mean voltage and electric field for each layer in both monkeys.

| | Monkey 1 | | Monkey 2 | |
|---|---|---|---|---|
| | Voltage (mV) | Electric field (V/m) | Voltage (mV) | Electric field (V/m) |
| Layer 1 | 0.35 | 0.62 | 0.17 | 0.97 |
| Layers 2/3 | 1.05 | 2.53 | 0.70 | 2.76 |
| Layer 4AB | 1.88 | 1.18 | 1.19 | 1.60 |
| Layer 4C | 2.12 | 0.62 | 1.44 | 1.59 |
| Layers 5/6 | 2.32 | 0.37 | 1.76 | 1.42 |
